# Stability of amino acids and related amines in human serum under different preprocessing and pre-storage conditions based on iTRAQ^®^-LC-MS/MS

**DOI:** 10.1101/2020.07.14.202473

**Authors:** Zhuoling An, Chen Shi, Pengfei Li, Lihong Liu

## Abstract

Amino acids analysis or metabonomics requires abundant serum/plasma samples collection and samples storage has become inevitable given the limited capacity for immediate analysis. Currently, most of the existing studies on metabolites stability during sample storage focused on long-term and short-term stability, while many functional amino acids might be ignored due to the poor sensitivity and detection of analysis methods. Here, we attempted to elucidate the stability of amino acids and related amines as comprehensive as possible in human serum following different preprocessing and pre-storage procedures. Pooled, fasting serum samples were collected and stored at 4 °C and 22 °C respectively after a delay in sample processing (0, 1, 2, 4, 8, 12 and 24 hours) and underwent freeze-thaw cycles for three times at −80 °C. The concentration of amino acids and related amines were quantified using isobaric tagging reagent iTRAQ^®^-LC-MS/MS. Approximately 54.84 %, 58.06 % and 48.39 % of detectable and target analytes altered at 4 °C and 22 °C during pre-treatment and freeze-thaw cycles. Some amino acids which are not stable and relatively stable were found. Our study provided detailed profiles and suggestions for amino acids in human serum corresponding to diverse collection and pre-treatment measures.

## Introduction

High-quality biological samples are required for reliable outcomes of research, as well as valid data sets, which are crucial for successful biomarker identification. The detection and quantification of free amino acids have been routinely applied in clinical laboratories for the diagnosis of inborn errors of metabolism [1]. The amino acid synthesis defects are presented by many clinical symptoms including central nervous system and mental disability for children, skin disorders such as cutis laxa in defects of proline synthesis, collodion-like skin and ichthyosis in serine deficiency, and necrolytic erythema in glutamine deficiency [2]. Increasing literatures have implicated the role of free amino acids in a number of diseases such as cardiovascular diseases [3-5], insulin resistance and type 2 diabetes [6,7], renal diseases [8], hepatic disorders [9] and cancer [10,11].

Increased interests in amino acids related to diseases have prompted the need for verifying the stability and sensitivity under different sample processing conditions. The existing studies almost all focused on long-term and short-term stability during sample storage. Indeed, storage of plasma samples at −80 °C for up to five years could lead to a change in concentrations of amino acids, acylcarnitines, glycerophospholipids, sphingomyelins and hexoses [12]. Augmented levels of amino acids during storage were observed in other long-term stability studies [13, 14]. The human plasma metabolome was adequately stable to long-term storage at −80 °C for up to seven years but the opposite result was found upon longer storage, where cysteine and cystine decreased over the analyzed storage period [15]. Evidence showed serum amino acids were modified in composition during 29 years long-term storage at −25 °C as methionine was transformed to methionine sulfoxide [16]. The stability of 18 free amino acids in the urine samples was found to be sustained for 72 hours at 4 °C, after one freeze thaw cycle but no more than four weeks at −80 °C [17]. The opposite results were found where amino acids were altered by delayed freezing [18]. The concentration of arginine, glycine, ornithine, phenylalanine, serine and isoleucine increased significantly during pre-storage handling at room temperature while glutamine decreased slightly at room temperature or on wet ice for 36 hours [19]. Stability in blood and plasma were also different due to the fact that platelets became activated and their metabolism was affected by low temperature [20].

Amino acids were markedly affected by preanalytical short-term storage in blood at room temperature for 2 hours and on wet ice for 6 hours, while it was prolonged to 16 hours in plasma at room temperature. The change in concentration of glutamate in plasma other than blood was observed presumably, while taurine changed only in blood through the same procedure [20]. Some amino acids and biogenic amines could become unstable within 3 hours on cool packs. Isoleucine, tryptophan and valine evidently decreased when undergone two freeze-thaw cycles [21]. The nine amino acids were stable up to 24 hours in plasma at 37 °C among which couple of amino acids changed statistically when the plasma specimens were placed at 4 °C for 24 hours [22]. The amino acids exhibited significant time-of-day variations [23]. Alanine and other metabolites changed after four or five freeze–thaw cycles at room temperature [24, 25].

Stability of amino acids and related amines in human serum varies under different preprocessing and pre-storage conditions, which is very important to our medical researchers because of the large number of serum/plasma samples collection for amino acids analysis in labs, which make it inevitable to optimize their storage as a result of the limited capacity for immediate analysis. However, a great deal of amino acids might be ignored and not detected due to the limitations of the scope and approach. The stability of amino acid metabolites have not been systematically studied in the past. Due to the lack of comprehensive research on amino acids, there is no comparison of multiple amino acids in the same dimension in one study, which leads to inconsistent or contradictory conclusions on the stability of some amino acids involved in different studies. Over the years, the cation-exchange and reversed phase liquid chromatography coupled to UV optical detection of pre-column or post-column derivatized have been processed on amino acids analysis. However, the co-eluting substances cannot be distinguished and quantified by these conventional approaches due to the lack of analyte specificity and selectivity. In the latest study, we have developed a novel approach by using stable isotope iTRAQ labeling and liquid chromatography tandem mass spectrometry to achieve comprehensive profiling and quantification of 42 amino acids and related amines. Compared with typical MS-based methods, the iTRAQ®-LC-MS/MS showed strong benefits in the availability of internal standards for all the analytes [26]. Our studies aimed to provide proof in estimating the stability of amino acids and related amines in serum samples after exposure adverse storage temperature and freeze-thawing cycles by using iTRAQ®-LC-MS/MS where the delineation of amino acid stability was submitted based on a comprehensive profiling and quantification approach of amino acids.

## Material and methods

### Chemicals and reagents

The derivatization of 44 amino acids and related amines were conducted on the iTRAQ^®^ Reagent Kit 200 Assay (P/N: A1116, AB Sciex, USA), including phosphoserine (PSer), phosphoethanolamine (PEtN), taurine (Tau), asparagine (Asn), serine (Ser), hydroxyproline (Hyp), glycine (Gly), glutamine (Gln), aspartate (Asp), ethanolamine (EtN), histidine (His), threonine (Thr), citrulline (Cit), sarcosine (Sar), *β*-alanine (*β*-Ala), alanine (Ala), glutamate (Glu), 1-methylhistidine, (1MHis), 3-methylhistidine (3MHis), argininosuccinic acid (Asa), carnosine (Car), homocitrulline (Hcit), arginine (Arg), *α*-aminoadipic acid (Aad), *γ*-aminobutyric acid (GABA), *β*-aminoisobutyric acid (*β*-Aib), *α*-amino-N-butyric acid (*α*-Abu), anserine (Ans), *δ*-hydroxylysine (*δ*-Hyl), proline (Pro), ornithine (Orn), cystathionine (Cth), cystine (Cys), lysine (Lys), methionine (Met), valine (Val), norvaline (Nva), tyrosine (Tyr), homocysteine (Hcy), isoleucine (Ile), leucine (Leu), norleucine (Nle), phenylalanine (Phe), tryptophan (Trp). Heptafluorobutyric acid (≥ 99.5%) was obtained from sigma Aldrich Corp (Switzerland) for mobile phase preparation. Acetonitrile and formic acid were purchased from Merck (Darmstadt, Germany). All chemicals and reagents were of appropriate analytical grades.

### Serum sample preparation under different storage conditions

The blood samples were collected from 11 different healthy volunteers (6 male, 5 female, age from 18 to 40 years) with overnight fasting status in Beijing Chao-Yang Hospital in September 2014. All participants were provided written informed consent. The study was carried out under the approval by the Ethics Committee of Beijing Chao-Yang Hospital affiliated with Beijing Capital Medical University and all experiments were performed in accordance with relevant guidelines and regulations. The whole blood sample collection was completed by professional medical staff. The patient information was verified and the collection time was recorded. About 4 ml of venous whole blood was collected from the cubital fossa vein of the upper limb of participants. The collected blood samples were placed in non anticoagulant tubes and allowed to coagulate at 4 °C for 10 minutes, then the samples underwent centrifugation at 3500 rpm at 4 °C for 10 minutes. The serum supernatant was separated and mixed in a new centrifugation tube by inverting, split into 200 *μ*L aliquots, and then subjected to three handling protocols: (1) the serum specimens were placed at 4 °C for 0, 1, 2, 4, 8, 12 and 24 hours; (2) samples were stored at 22 °C for 0, 1, 2, 4, 8, 12 and 24 hours; (3) serum aliquots were stored at −80°C and subjected to up to three freeze-thaw cycles. A freeze-thaw cycle consisted of taking out from −80 °C, thawing aliquots for one hour at 4 °C, and setting them back to −80 °C for 12 hours. Experimental design for stability of amino acids and related amines in human serum under different storage conditions were showed in Fig.1. Serum specimens were immediately frozen in liquid nitrogen to stop reaction and preserved respectively.

**Fig. 1.**
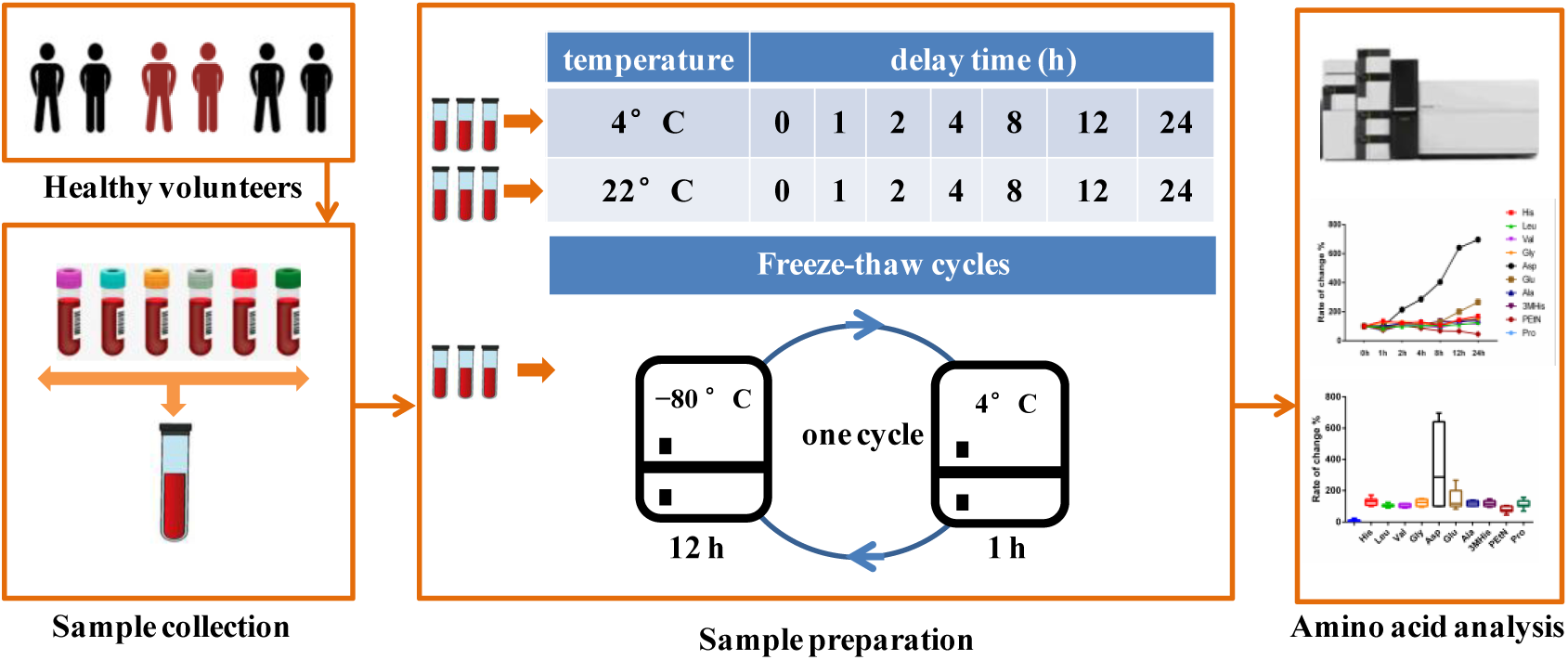
Scheme of stability investigation of amino acids and related amines in human serum under three preprocessing and pre-storage. (1) the serum specimens placed at 4 °C for 0, 1, 2, 4, 8, 12 and 24 hours; (2) samples were placed at 22 °C for 0, 1, 2, 4, 8, 12 and 24 hours; (3) serum aliquots were stored at −80°C and subjected to up to three freeze-thaw cycles. A freeze-thaw cycle consisted of taking out from −80 °C, thawing aliquots for one hour at 4 °C, setting them back to −80 °C for 12 hours.

### Targeted metabolite quantification

Each sample was measured according to the procedures reported in our previous method [26]. Quadruplicate samples were processed and tested parallel at same temperatures and conditions for equal duration. A workflow indicating the procedures for biological sample preparation and amino acid derivatization using ITRAQ reagents was shown in Fig.2. An aliquot of 40 *μ*L of serum samples was supplemented with 10 *μ*L of 10% sulfosalicylic acid containing 4 nmol of norleucine. The norleucine was applied as IS for the evaluation of extraction efficiency. The mixture underwent vortex for 30 s and centrifuged at 10,000 *g* for 2 min at 4 °C. 10 *μ*L supernatant was added with 40 *μ*L labeling buffer (containing 20 *μ*mol/L norvaline for evaluation of derivatization efficiency). Subsequently, 10 *μ*L of the diluted supernatant after vortex for 30 s was mixed with 5 *μ*L iTRAQ^®^ reagent 121 solution (one tube mixed with 70 *μ*L isopropanol). The termination of derivatization reaction was implemented by adding 5 *μ*L of 1.2 % hydroxylamine solution to the mixture after incubating at room temperature for 30 min. The resulting mixture was evaporated to dryness under a nitrogen stream. Finally, we re-dissolved the dried residue with 32 *μ*L iTRAQ^®^ reagent 113-labeled standard mix [26].

**Fig. 2.**
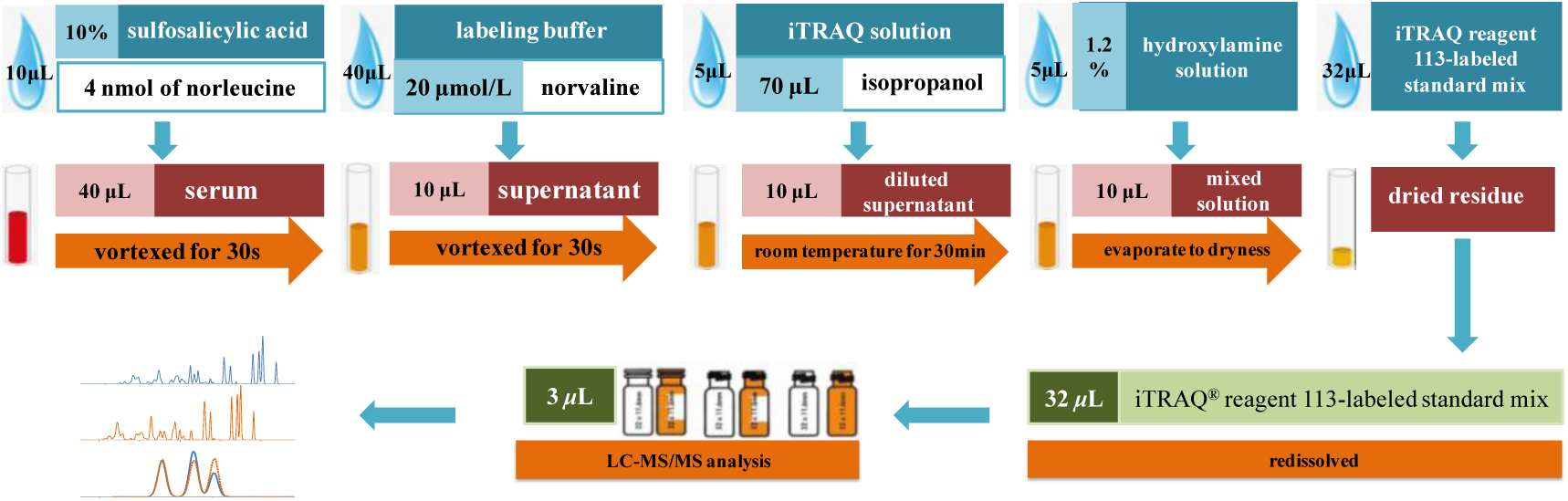
A workflow indicating the procedures for biological sample preparation and amino acid derivatization using ITRAQ reagents.

### Instrumental analysis

The analysis of amino acid derivatives was performed on a Shimadzu LC-20AT liquid chromatography system couple with an API 3200 Qtrap™ mass spectrometer. Chromatography was performed on a XBridge Shield RP18 column (5 *μ*m, 150 mm×4.6 mm) where the column temperature was held constant at 50 °C with the injection volume of 3 *μ*L. Mobile phase A was water and phase B was acetonitrile, both containing 0.01% heptafluorobutyric acid and 0.1% formic acid at a flow of 0.8 mL/min. The separation was conducted under the following gradient: 0–11 min, 0–20 % B; 11–11.5 min, 20–100 % B; 11.5–14 min, 100 % B; 14–14.1 min, 100–0 % B; 14.1–21 min, 0 % B. Multiple reaction monitoring (MRM) in positive ionization mode was used for amino acid derivatives detection. Parameters including ESI voltage, entrance potential (EP), and declustering potential (DP) was set up to +5.5 kV, 30 V and 10 V respectively. Besides, 540 °C source temperature, 20 psi curtain gas flow was, 50 psi nebulizer gas flow was, 60 psi source gas flow and 30 eV collision energy (CE) were displaced in this study. Data acquisition was carried out by Analyst 1.5.1 software on a DELL computer.

### Data processing and statistical analysis

Peak integration of iTRAQ-121 and iTRAQ-113 labeled amino acids were carried out by the Analyst 1.5.1 software (Applied Biosystems Sciex). The iTRAQ-113 labeled amino acids were used as the isotopic ISs for the normalization of their corresponding iTRAQ-121 labeled amino acids in biological samples. The obtained quantitative data of amino acid were further used for statistical analysis. SIMICA-P+ 13.0 (Umetrics AB, Umeå, Sweden) was employed for the principal component analysis (PCA) analysis. The changes of amino acid levels affected by various environmental factors were identified by using t-test (*p*<0.05).

## Results

### Quantification of amino acids and related amines by iTRAQ-based profiling

A novel approach had been developed for the comprehensive profiling and quantification of amino acids and related amines based on iTRAQ^®^-LC-MS/MS in our previous study [26]. Norvaline was added to the reaction system to investigate the derivatization efficiency. Results showed more than 80% norvaline could be derivatized by the iTRAQ reagent. Thirty-one labeled amino acids and related compounds were separated and quantified with excellent peak shapes in current study. The MRM ion chromatograms corresponding to amino acids and their isotopic internal standards were extracted from the data obtained by LC-MS/MS. Integration and calculation were adjusted to quantify amino acid levels to explore the effects of pre-processing temperature and freeze-thawing cycles upon the stability of amino acids and related amines in serum samples. Principal component analysis (PCA) could provide the information of clustering in each group and the possible changes of the metabolic profiles. The concentration was assigned as variable while the pre-storage handing conditions and freeze-thaw cycles were set as factors in the multivariate data of quantification results of 31 amino acids. SIMCA-P 13.0 (VersionAB, Umeå, Sweden) were employed to visually investigate the clustering of amino acids detectable in serum at different pre-processing and conditions. Fig.3 showed the three-dimensional PCA score plot of serum samples placed at 4 °C and 22 °C with different dosage duration as well as samples detected immediately after processing process. This multivariate analysis showed relative values of R2X (cum) = 0.385 and Q2 (cum) = 0.203. The samples placed at 4 °C and 22 °C were clustered obviously and the samples stored at 22 °C were relatively scattered and more deviation were found in samples immediately analyzed after processing. Fig.4 showed the three-dimensional PCA score plot of serum samples for different dosage duration (0, 1, 2, 4, 8, 12 and 24 hours) at 4 °C (A) and 22 °C (B) respectively. The values for the two multivariate analysis were R2X (cum) = 0.427, Q2 (cum) = 0.185 and R2X (cum) = 0.442, Q2 (cum) = 0.213, respectively. Moreover, amino acids had been changed in a time-dependent manner at different pre-processing temperature. The serum samples at 4 °C from 0 to 24 hours scattered and distributed with prolonged storage time, gathered at the same pre-storage times but gradually deviated from immediately detected samples. While placed at the room temperature, the serum samples scattered and distributed relying on the time extension, however they deviated from the sample group with rapid detection significantly during 8 to 24 hours. Taken together, the results showed that temperature and sample store time had a greater impact on the composition of amino acids in serum.

**Fig. 3.**
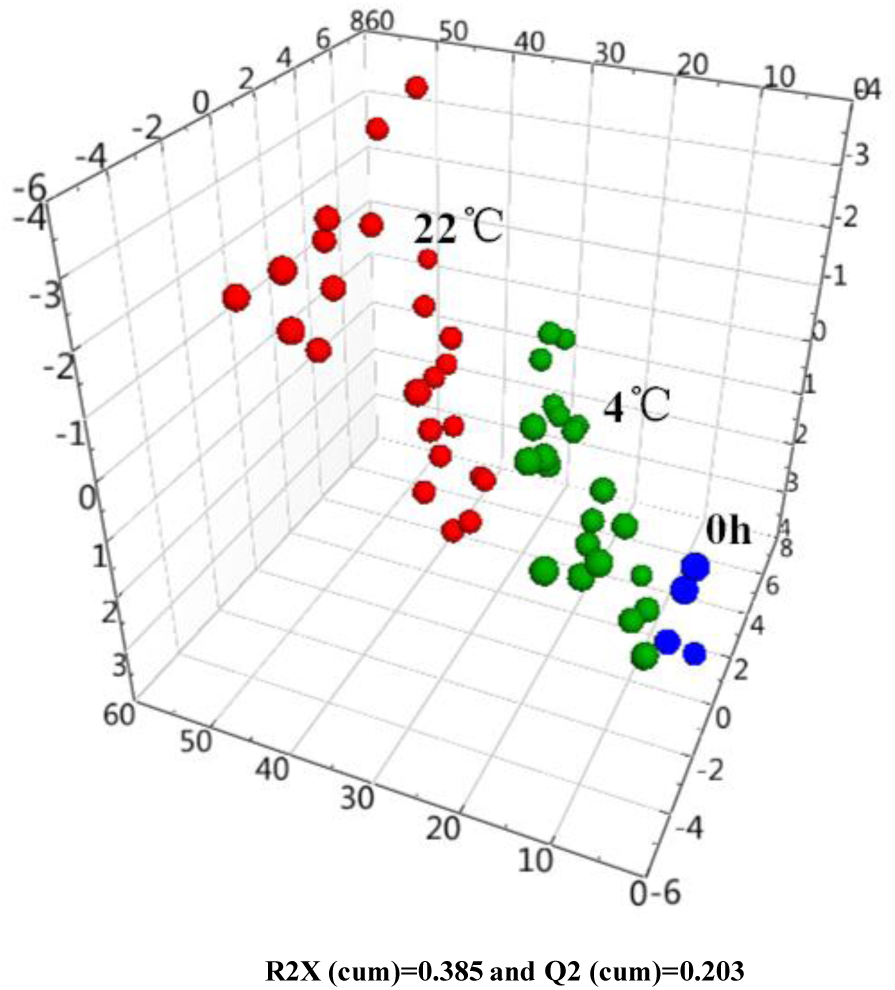
PCA score plot of the concentration of amino acids in human serum deposited at 4 °C (**◍**), 22°C (**◍**) and samples detected immediately after processing process (**◍**).

**Fig. 4.**
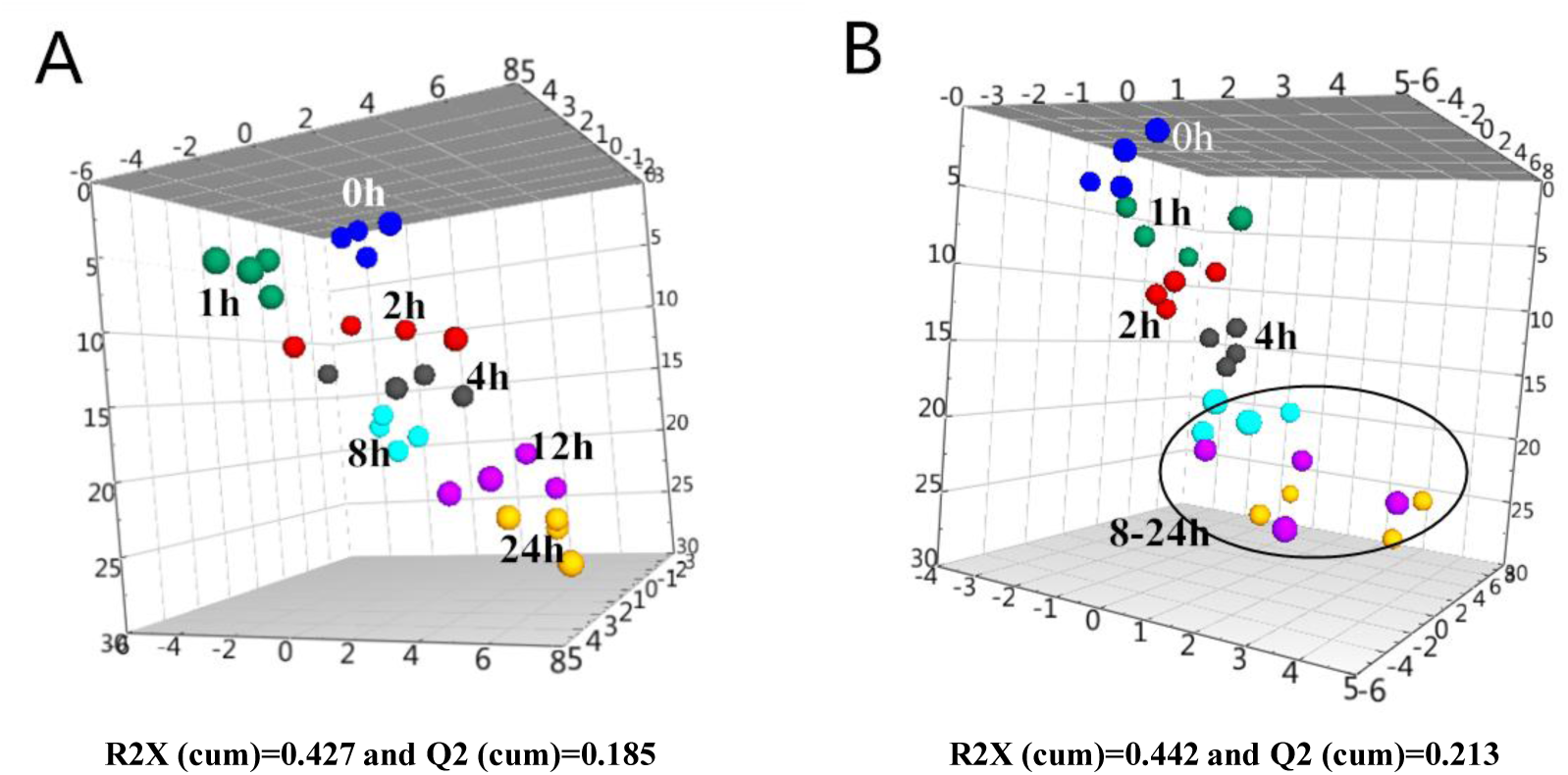
PCA score plot of the serum samples deposited at 4 °C (A) and 22 °C (B) from 0 h to 24 h (**◍**:0 hour,**◍**:1 hour,**◍**:2 hours,**◍**:4 hours,**◍**:8 hours,**◍**:12 hours,**◍**:24 hours).

### Stability of serum amino acids at 4 °C

17 amino acids in serum were significantly altered at 4 °C with different pre-processing periods (*p*<0.05) (Fig.5). The content and change rates of these amino acids were altered significantly after serum specimens incubated at 4 °C for 24 hours. (Table S1). Histidine, leucine, phenylalanine, tryptophan, valine, glycine, aspartate, glutamate, *β*-alanine and 3-methylhistidine increased significantly after 24 hours. Lysine and taurine were differentially regulated to significantly higher levels within 4 hours while serine, cystine and alanine decreased after 8 hours and phosphoethanolamine as well as proline decreased after 24 hours. It should be noted that aspartate remained relatively stable within one hour, nonetheless the concentration increased rapidly after one hour to approximately seven hundred-fold changes. Glutamate also increased continuously after 8 hours with the change rate of 266 %.

**Fig. 5.**
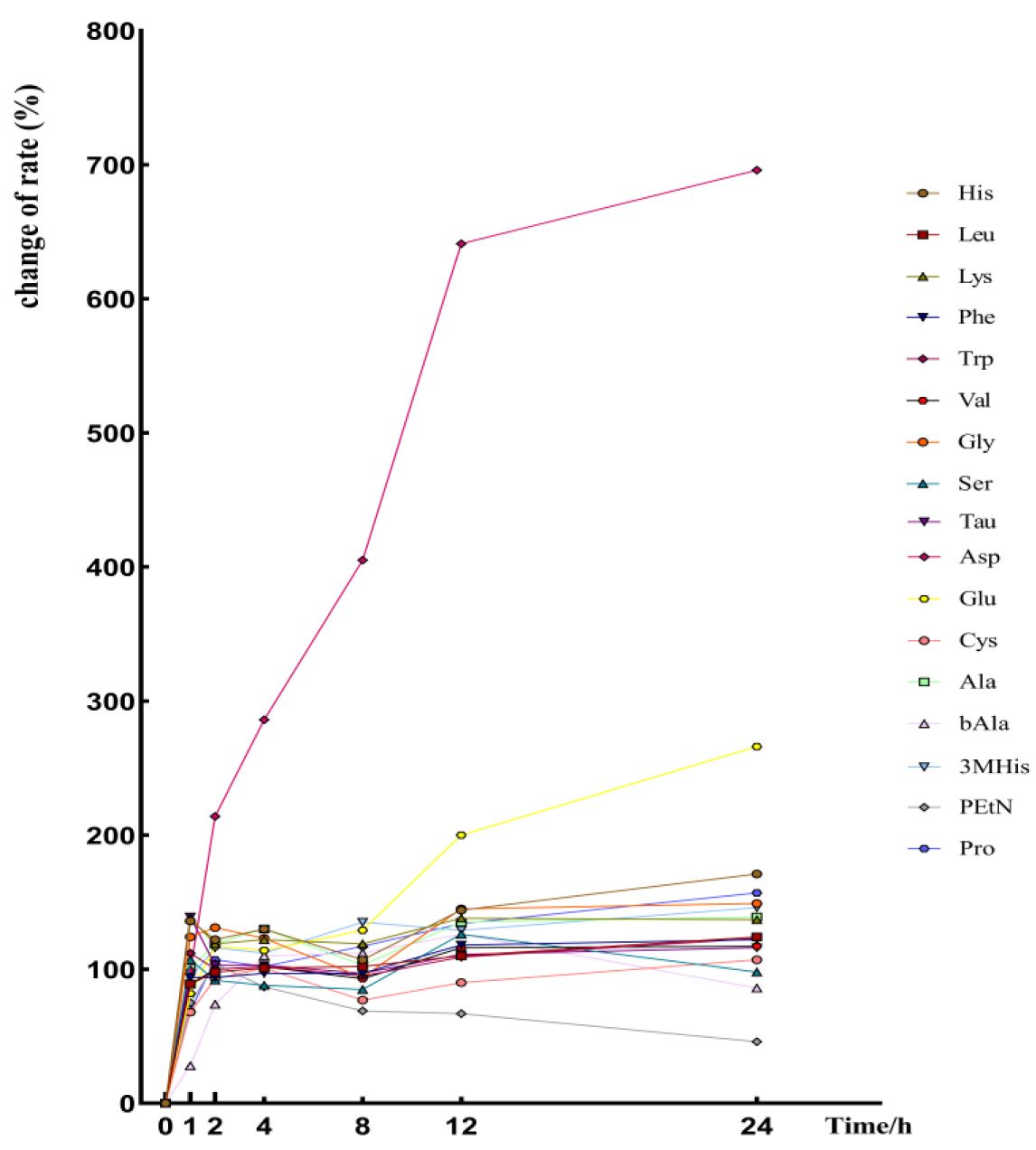
The change rate of 17 amino acids in serum after serum specimens incubated at 4 °C for 24 hours (*p*<0.05).

### Stability of serum amino acids at 22 °C

The levels of 18 amino acids markedly varied after 24 hours at room temperature after serum sample preprocessing (*p*<0.05) (Fig.6). Histidine, isoleucine, leucine, lysine, methionine, phenylalanine, valine, aspartate, glycine, ornithine and 3-methylhistidine increased obviously after 24 hours (Table S2). A reduction of cystine and phosphoethanolamine occurred after 24 hours of storage as threonine and tryptophan decreased within 12 hours. The levels of asparagine, glutamate and alanine were markedly elevated after 4 hours. Moreover, approximately three hundred-fold changes of the aspartate occurred rapidly within one hour and became concentrated steadily thereafter. The change rate of aspartate had reached almost 634% until 24 hours. Glutamate continuously increased significantly after two hours of preprocessing and the change rate was about 462 %. Cystine and phosphoethanolamine decreased about 16% to 47% compared with their initial concentration after 24 hours of storage.

**Fig. 6.**
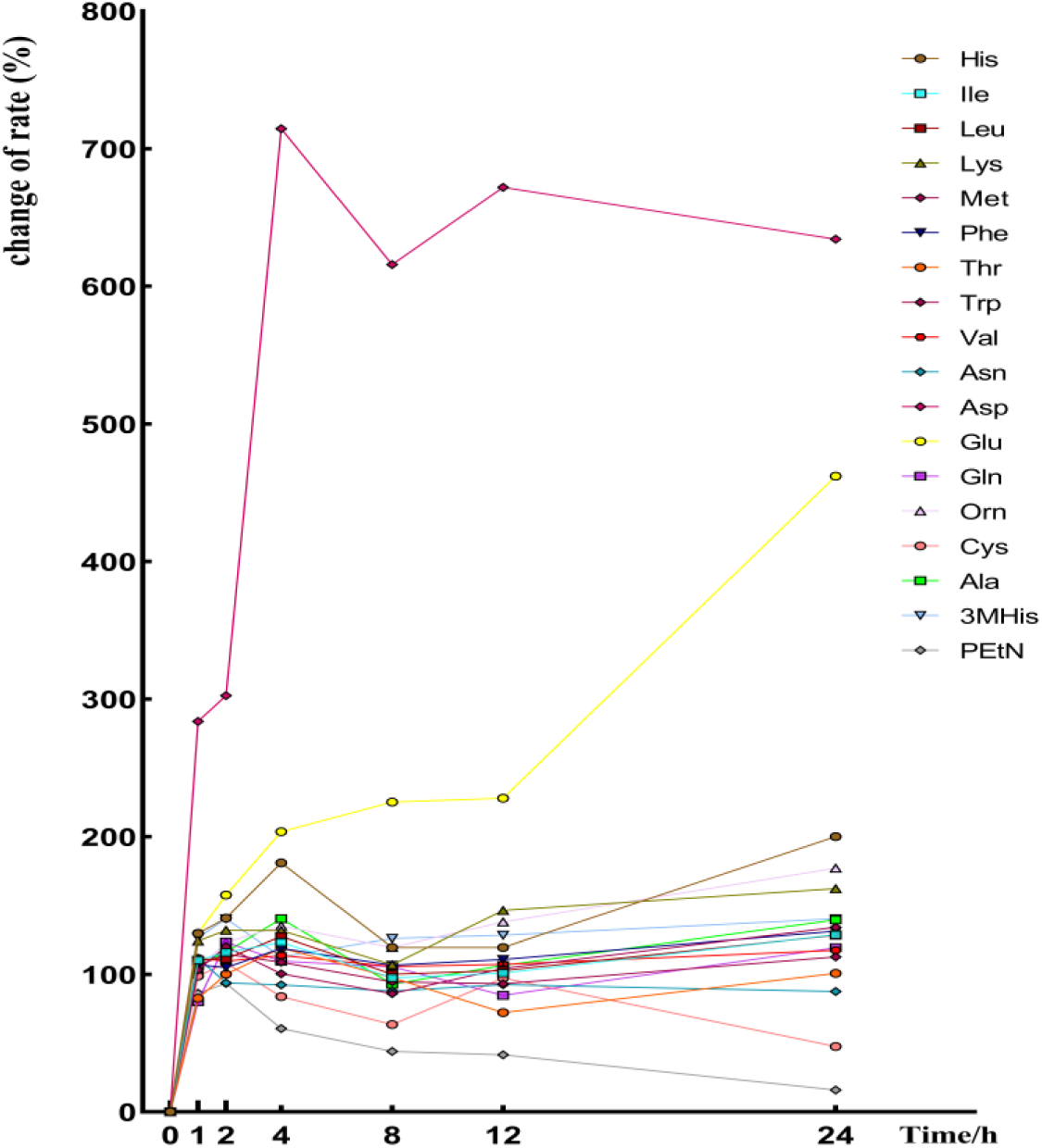
The change rate of 18 amino acids in serum after serum specimens incubated at 22 °C for 24 hours (*p*<0.05).

### Stability of serum amino acids after three freeze-thaw cycles at −80 °C

After the serum samples were frozen and thawed three times at −80 °C, the concentrations of 11 amino acids (histidine, leucine, isoleucine, methionine, phenylalanine, glutamate, tryptophan, valine, taurine, tyrosine and ornithine) in the serum increased obviously compared with the samples without freeze-thaw treatment (*p*<0.05). Four amino aicds including cystine, *β*-alanine, 1-methylhistidine and aspartate reduced in concentration after the freeze-thaw cycle (*p*<0.05) (Fig.7). The increased rate of change for these amino acids varied from 31.45 % to 252.12 % while that of the reduced rate was 42.57 % to 100.00 % (Table S3).

**Fig. 7.**
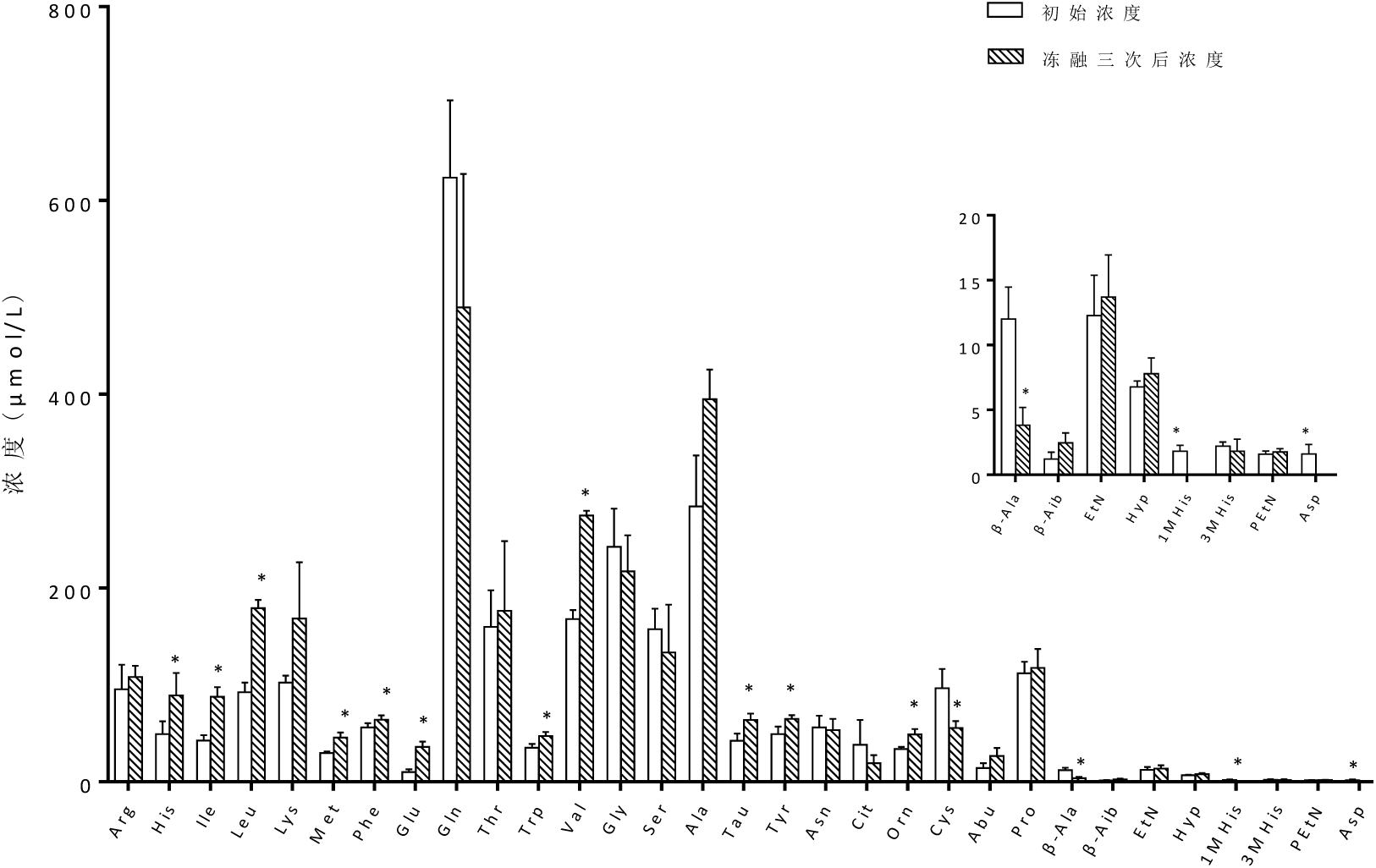
The changes of amino acids in serum after three freeze-thaw cycles and the concentrations of 14 amino acids in the serum altered obviously compared with the samples without freeze-thaw process (*p*<0.05). The freeze-thaw cycle including samples were brought from −80 °C, placed at 4 °C for one hour an delivered to −80 °C for 12 hours.

### Marked concentration changes in amino acids due to pre-storage handling

Our experiment with different pre-storage handling conditions showed pronounced changes of the detectable amino acids and related amines. Approximately 54.84 %, 58.06 % and 48.39 % of target analytes altered at 4 °C and 22 °C and freeze-thaw cycles during preprocessing. Seven amino acids, regarding to histidine, leucine, phenylalanine, tryptophan, valine, aspartate and cystine,were more sensitive with evident modifications by multiple factors including different storage time and temperature as well as repeated freeze-thaw cycles. On the contrary, arginine, tyrosine, citrulline, *α*-amino-N-butyric acid, *β*-aminoisobutyric acid, ethanolamine and hydroxyproline were more stable and remained unchanged under any conditions. Except tyrosine, all branched chain amino acids and aromatic amino acids were sensitive to storage temperature and freeze-thaw. It was noteworthy that the augmented levels of aspartate and glutamate were observed during 4 °C and 22 °C for extended storage periods in our study. This possibly results from that asparagine and glutamine were converted to their dicarboxylic acid counterparts by deamidation [27]. Protein degradation process is the key factor for the increase of amino acids, especially for those amino acids occurring with high frequency in proteins e.g. isoleucine and glycine. Tryptophan and phenylalanine increased distinctly after repeated freeze-thaw cycles given the protein degradation during thawing and refreezing. This assumption was supported by the fact in previous study [19, 28]. Increased cystine could be explained by rapid oxidation from instable cysteine to cystine at room temperature. However, since level of cystine was also lowered during repeated freeze-thaw cycles, the reduction of both cysteine and cystine could be a result of oxidative conversion to unidentified derivates as described previously [29]. Up to four freeze-thaw cycles will not affect the stability of the metabolites which was inferred by other researchers [19]. Conversely, our results indicated the freeze-thaw had a great effect on the stability of the amino acids in biological samples.

## Conclusions

The stability of amino acids in serum samples which underwent prolonged exposure at 4 °C and 20 °C and repeated freeze-thaw cycles at −80 °C were investigated by using stable isotope iTRAQ labeling and liquid chromatography tandem mass spectrometry. The results indicated that pronounced changes of amino acids had occurred during different preprocessing and the pre-storage conditions separately. Temperature, storage time and freeze-thaw cycle imposed critical effects on the stability of amino acids. In that case, the precise control over the collection, preservation and pretreatment of serum samples should be optimized and standardized in certain aspects, for example, biological samples should avoid freeze-thaw before detection. Meanwhile, the process of pretreatment should be operating at 4 °C to improve the reliability of potential biomarkers and metabolites. The current study provided a standard method for the collection, preparation, transportation and storage of serum samples through the quantitative analysis of amino acids and metabonomics based on liquid chromatography-mass spectrometry.

